# Stability of *β*-lactam antibiotics in bacterial growth media

**DOI:** 10.1101/2020.04.15.044123

**Authors:** Rebecca Brouwers, Hugh Vass, Angela Dawson, Tracey Squires, Sharareh Tavaddod, Rosalind J. Allen

## Abstract

Laboratory assays such as MIC tests assume that antibiotic molecules are stable in the chosen growth medium - but rapid degradation has been observed for antibiotics including *β*-lactams under some conditions in aqueous solution. Degradation rates in bacterial growth medium are less well known. Here, we develop a ‘delay time bioassay’ that provides a simple way to estimate antibiotic stability in bacterial growth media. We use the bioassay to measure degradation half-lives of the *β*-lactam antibiotics mecillinam, aztreonam and cefotaxime in widely-used bacterial growth media based on MOPS and Luria-Bertani (LB) broth. We find that mecillinam degradation can occur rapidly, with a half-life as short as 2 hours in MOPS medium at 37°C and pH 7.4, and 4-5 hours in LB, but that adjusting the pH and temperature can increase its stability to a half-life around 6 hours without excessively perturbing growth. Aztreonam and cefotaxime were found to have half-lives longer than 6 hours in MOPS medium at 37°C and pH 7.4, but still shorter than the timescale of a typical minimum inhibitory concentration (MIC) assay. Taken together, our results suggest that care is needed in interpreting MIC tests and other laboratory growth assays for *β*-lactam antibiotics, since there may be significant degradation of the antibiotic during the assay.

## Introduction

Antibiotic efficacy is usually quantified by the minimal inhibitory concentration (MIC), or the concentration of antibiotic needed to prevent bacterial growth over 24 hours in a standard laboratory growth medium [1]; the MIC value plays a central role in diagnostic and therapeutic decision-making. When performing an MIC assay, one assumes that the antibiotic does not degrade over the timescale of the assay. Antibiotic stability is also assumed in a host of other bacterial growth assays that are used to determine antibiotic mechanism of action [2–5], interactions between antibiotics [6–9], and evolution of antibiotic resistance [10–14].

Antibiotic degradation in aqueous solution has been extensively studied in the chemical literature [15–19], and it is well-known that, under some conditions, antibiotics can degrade on timescales much shorter than a typical bacterial growth assay. There has been little work, however, on how quickly antibiotics degrade in standard laboratory growth media, such as those used for MIC assays. Here, we develop a ‘delay-time bioassay’, based on growth measurements in a plate reader, that allows simple estimation of the rate of antibiotic degradation in bacterial growth media.

Bacterial growth media are chemically complex, since bacterial proliferation requires carbon, nitrogen, phosphorous, iron and a diverse array of micronutrients [20, 21]. “Undefined” growth media contain ingredients such as yeast extract, blood or beef extract, whose detailed chemical composition is not known. Luria-Bertani (LB) medium, which consists of water, tryptone, sodium chloride and yeast extract, is a widely-used example. In contrast, “defined” growth media contain known quantities of defined chemical ingredients, and are often pH-buffered, typically at pH 7.2 ± 0.2. For example, Neidhardt’s rich defined medium [22] consists of a MOPS (3-(N-Morpholino)propane sulfonic acid) pH buffer supplemented with glucose, amino acids, nucleotides and sources of sodium, potassium, chlorine, phosphorous, magnesium, iron, zinc, manganese, bromine, boron, copper, cobalt and molybdenum. It is reasonable to expect that microbiological growth media might affect antibiotic degradation rates; for example, transition metals have been reported to accelerate degradation of *β*-lactam antibiotics [23], and Neidhardt’s medium contains Fe^2+^ at 10 *µ*M, as well as Cu^2+^ at 10 nM, Mn^2+^ at 80 nM and Co^2+^ at 30 nM [22]. Although we are not aware of any quantitative measurements of degradation rates in bacterial growth media, several studies have commented on the phenomenon of ‘regrowth’, in which bacterial cultures show late-time growth in *β*-lactam-supplemented media after apparent early-time growth suppression [24–27]. This phenomenon could arise from either resistance evolution or degradation of the antibiotic.

Our focus here is on degradation rates for *β*-lactam antibiotics. *β*-lactams, which inhibit bacterial cell-wall synthesis [28–30], account for the majority of global antibiotic consumption [31]. They are characterised by a *β*-lactam ring in their molecular structures (Fig 1); clinically relevant classes of *β*-lactams include penicillin derivatives, cephalosporins, monobactams and carbapenems. The degradation of *β*-lactam antibiotics in aqueous solution is known to be strongly pH-dependent [32]; *β*-lactams are degraded via different pathways under acidic versus basic conditions, typically having a U-shaped profile for degradation rate as a function of pH, with maximal stability around pH 4-5 for *α*-amino *β*-lactams (e.g. amoxicillin and ampicillin) and around pH 6-7 for *β*-lactams without a side-chain amino group [33]. As noted above, the degradation rate can also be very sensitive to the presence of transition metal ions [23, 32, 33].

**Fig 1.**
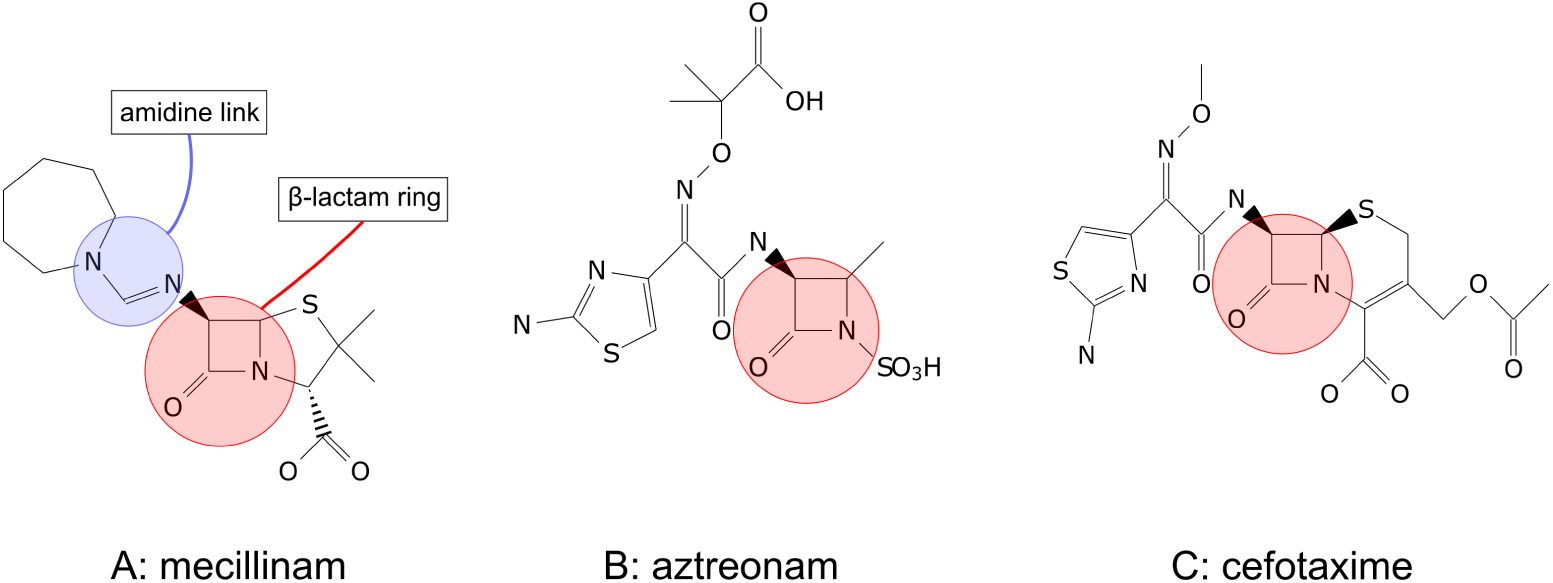
Chemical structures of mecillinam, aztreonam and cefotaxime. A: Mecillinam has a characteristic penicillin structure with a fused beta-lactam-thiazolidine two-ring system [39] and an amidine link to the side chain. B: Aztreonam has a monobactam structure with no secondary ring attached to the *β*-lactam ring, and with an N-SO_3_H side-chain [35]. C: Cefotaxime has a characteristic cephalosporin structure with a fused *β*-lactam-Δ^3^-dihydrothiazine two-ring system [39]. All structures were drawn using the chemfig Latex package.

Our study focuses on three *β*-lactams: mecillinam, aztreonam and cefotaxime (Fig 1) which have previously been shown to have stability maxima at pH 4-6 [34–36]. Mecillinam (also known as amdinocillin or FL1060) is an amidinopenicillin that binds selectively to the PBP2 transpeptidase, inhibiting peptidoglycan synthesis during the elongation phase of bacterial growth. Mecillinam is active against Gram-negative bacteria, especially Enterobacteriaciae, and is mainly used to treat urinary tract infections. Mecillinam is somewhat peculiar among *β*-lactams since its side chain is linked to the core part of the penicillin molecule (the 6-aminopenicillanic acid moiety) via an amidine bond rather than the more usual amide bond (Fig 1A). Its degradation has been shown to be highly pH-sensitive in aqueous solution, with a maximum half-life at pH 5 (at 37°C) of around 200 hours [16]. Aztreonam (Fig 1B) is a synthetic monobactam that targets peptidoglycan synthesis during cell division by binding to the PBP3 component of the peptidoglycan-synthesizing divisome complex [37]. Aztreonam is only active against Gram-negative bacteria. It has been found to be most stable at approximately pH 5 in aqueous solution (at 35°C) [35], and to be more stable than other *β*-lactams in neutral or acidic conditions. The increased stability is possibly due to reduced strain on the *β*-lactam ring as it is not attached to a secondary ring (Fig 1B) [35]. Cefotaxime (Fig 1C) is a third-generation cephalosporin *β*-lactam that targets multiple components of the peptidoglycan synthesis machinery, but with a high affinity for the PBP3 transpeptidase and the bifunctional TPase/TGase PBP1b [38], which are involved in cell division [37]. Cefotaxime is active against a wide range of bacterial species [39] and is used to treat meningitis and septicaemia [40]. The rate of degradation of cefotaxime has been found to be strongly affected by solvolytic, hydrogen ion and hydroxide ion catalysis, with maximum stability at pH 4.5 - 6.5 (at 25°C) in aqueous solution [36].

Our results show that mecillinam is rather unstable in both MOPS-based media and in LB (S3 Fig), but that (for MOPS media) its stability can be enhanced by adjusting the pH and temperature without excessively compromising bacterial growth. Aztreonam and cefotaxime were found to be more stable, but with half-lives that are still much shorter than the timescale of a typical MIC assay. Therefore our work suggests that care is needed in interpreting MIC tests and other laboratory growth assays for *β*-lactam antibiotics, since there may be significant degradation of the antibiotic during the assay.

## Results

### Growth assays under standard conditions show regrowth

Bacteria exposed to *β*-lactam antibiotics can sometimes show “regrowth”: bacterial growth is initially suppressed by the antibiotic, but at late times the culture starts to grow, eventually reaching a significant cell density [24–27]. Regrowth could be due to degradation of the antibiotic or to the emergence of resistance in the bacterial population. We observed consistent regrowth in plate reader growth assays for *E. coli* (strain RJA002, an isogenic fluorescent derivative of MG1655; see Methods) in the presence of mecillinam, aztreonam and cefotaxime (Fig 2 and S3 Fig; see also Methods). After a long period of growth inhibition, the bacterial population started to grow at late times, reaching a significant density by the end of our 24 hour experiments (Fig 2). The length of time before regrowth happened generally increased with the antibiotic concentration (Fig 2). Regrowth was observed on LB medium (S3 Fig C), on MOPS rich defined medium with glucose (MOPSgluRDM; see Methods), and on other MOPS-based media supplemented with glucose or glycerol, in the presence or absence of amino acids and nucleotides (Fig 2 and S3 Fig D-F).

**Fig 2.**
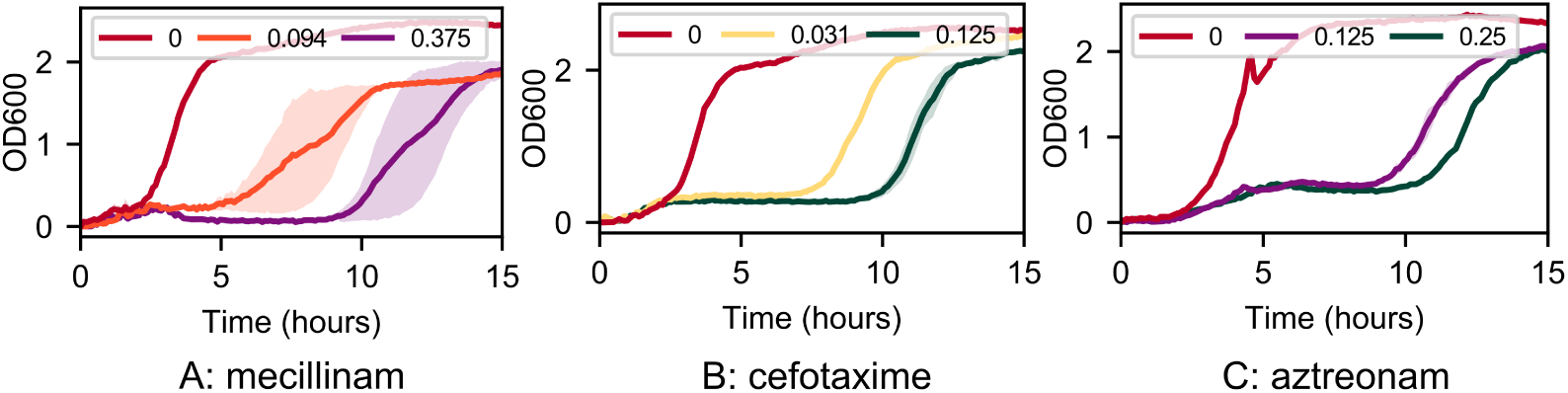
Typical growth curves in the presence of *β*-lactam antibiotics. Growth curves, measured in the plate reader, for *E. coli* strain RJA002 (see Methods) in MOPSgluRDM, in the presence of A: mecillinam, B: cefotaxime and C: aztreonam. The legends show the antibiotic concentration, in units of *µ*g/ml. Growth curves are averaged over two replicates. The standard deviation between the replicates is represented by the shaded area around the lines.

### A delay-time assay shows rapid degradation of mecillinam in MOPSgluRDM

Motivated by our observations of regrowth in bacterial growth assays, we designed a “delay time bioassay” to measure antibiotic degradation in bacterial growth media. This assay uses a 96-well microplate setup in which replicate wells containing antibiotic are inoculated with bacteria at different times after the start of the experiment. The setup is shown in detail in Fig 3; antibiotic dilutions are set up identically in replicate columns, but different columns are inoculated with *E. coli* at different times (see Methods for further details). Bacterial growth is monitored throughout the experiment via OD measurements in a plate reader. If the antibiotic is not degraded, then we would expect to see identical sets of growth curves in replicate columns, simply shifted by the inoculation time. However, if the antibiotic is degraded, we expect to observe different growth curves depending on the inoculation time (since the effective concentration of antibiotic at the time of inoculation will be lower for later inoculations). Fig 4 (a) shows that the latter scenario is indeed observed, for *E. coli* growing in MOPSgluRDM with mecillinam. Here, the dashed lines correspond to wells inoculated 2 hours later than the solid lines, while the colours indicate antibiotic concentration. For example, a comparison between the orange dashed and solid lines shows that growth at the initial concentration of 0.094 *µ*g/ml mecillinam is considerably enhanced for the later inoculation.

**Fig 3.**
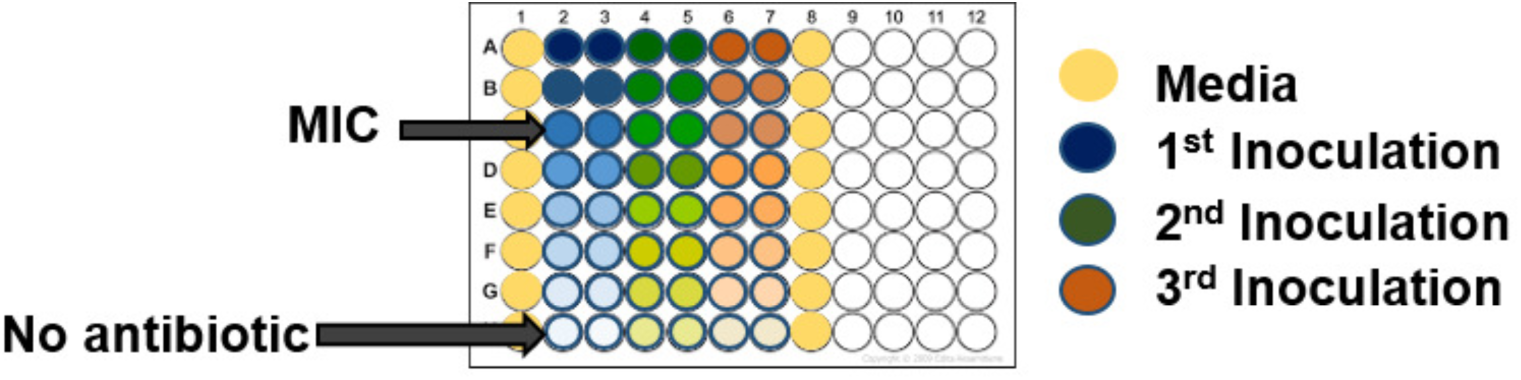
Plate setup for the delay-time bioassay. The entire plate is filled with media + antibiotics at the start of the experiment, with dilution series down the columns as indicated by the shading. The wells shown in yellow are media-only controls. The blue set of replicate columns are inoculated with bacteria immediately, and the plate is then incubated in a plate reader with OD measurement. The green set of columns are inoculated 2 hours later, the pink set of columns a further 2 hours later, etc. If the antibiotic has degraded between inoculations, this will be reflected in a change in the shape of the measured growth curves for later inoculations.

**Fig 4.**
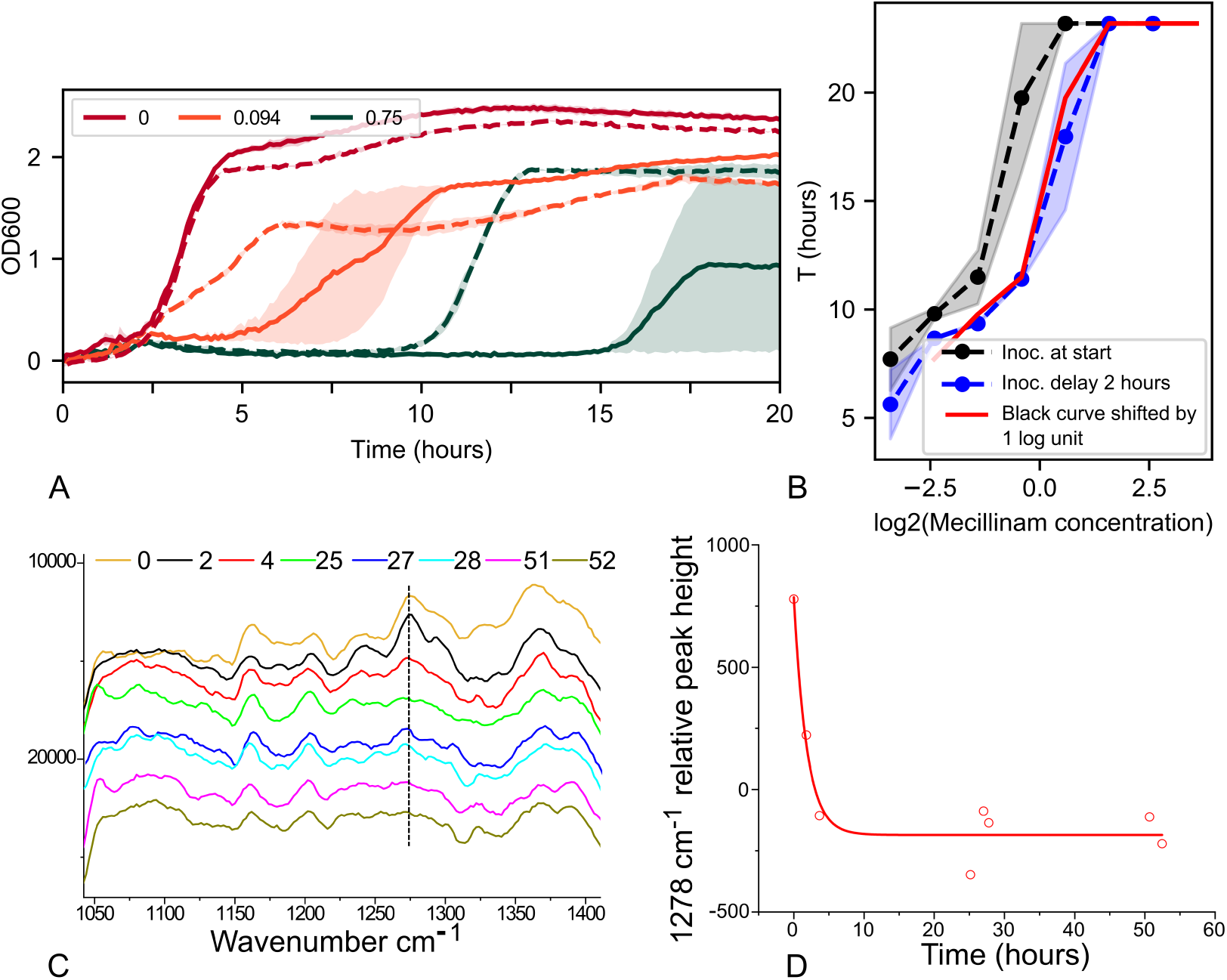
Degradation half-life of mecillinam is approximately 2 hours in MOPSgluRDM at pH 7.4, 37°C. A: Bioassay measurements: growth curves for *E. coli* strain RJA002 in MOPSgluRDM at pH 7.4, 37°C, for cultures inoculated at time zero (solid lines) and after a delay period of 2 hours (dashed lines). The different colours correspond to different (initial) concentrations of mecillinam, as displayed in the legend, in *µ*g/ml. Only a selection of the concentrations used are displayed to improve the clarity of the plot. B: Bioassay analysis - the time *T* at which the growth curves reach a threshold OD of 0.75 is plotted as a function of the initial antibiotic concentration. The black data correspond to inoculation at time zero; the blue data to inoculation after a 2 hour delay. The red line shows the black data, shifted by one unit on the log_2_ scale; since the blue data falls onto the red line, the degradation half-life is approximately 2 hours. In both panels A and B, the data is averaged over two replicates and the shaded areas represent the standard error. C,D: Raman spectroscopy confirms rapid degradation of mecillinam in MOPSgluRDM. C: Raman spectra for the region between 1000 and 1400 wavenumbers, recorded at times 0 to 52 hours (time in hours indicated by colour as shown in the legend). Successive curves are shifted upwards for clarity. D: Relative peak height at 1278 cm^−1^ as a function of time is fit to an exponential decay, which gives a half-life of 1.3±0.4 hours. The peak height at 1278 cm^−1^ was measured relative to the background spectrum (see Methods).

The delay-time bioassay can be used to estimate the degradation half-life of the antibiotic in the chosen growth medium. For a culture that is inoculated at time *t*_*ic*_, the effective antibiotic concentration at the inoculation time will be *a*_*eff*_ = *a*_0_ exp (− ln 2 × *t*_*ic*_*/t*_*d*_) where *a*_0_ is the antibiotic concentration supplied at the start of the experiment and *t*_*d*_ is the degradation half-life of the antibiotic. We therefore expect the measured growth curve in that well to match the growth curve measured for a well containing concentration *a*_*eff*_, inoculated at time zero. Specifically, for a well set up with a given antibiotic concentration and inoculated at a time that equals the degradation half-life, *t*_*ic*_ = *t*_*d*_, the measured growth curve should match that of a well that was set up with half the concentration, but inoculated at time zero.

To simplify our analysis, we chose to characterise the growth curves by a single number: the time *T* after inoculation at which the OD reaches a threshold value OD_*t*_ (here taken to be OD_*t*_ = 0.75). Because the antibiotic inhibits growth, the threshold OD value is reached later for higher antibiotic concentrations. Thus the value of *T* increases with the effective antibiotic concentration. Plotting *T* versus the logarithm of the antibiotic concentration (for a fixed inoculation time), we obtain an increasing curve (Fig 4 B; black curve is for inoculation at time zero). Repeating this plot for a later inoculation time, we observe a shift of the entire curve to the right, because the effective antibiotic concentration has decreased during the inoculation delay (Fig 4 B; blue curve is for inoculation at time *t*_*ic*_ = 2 hours). If the antibiotic concentration axis is plotted on a log_2_ axis, then an inoculation time equal to the degradation half-life (*t*_*ic*_ = *t*_*d*_) results in a shift of the curve by one unit along the concentration axis. In Fig 4 B, the red curve shows the black data shifted by one unit; if the blue curve matches the red curve, the inoculation time equals the degradation half-life. For mecillinam in MOPSgluRDM at pH 7.4 and 37°C this is the case for an inoculation time of 2 hours (Fig 4 B); therefore the degradation half-life is approximately 2 hours.

### Raman spectroscopy confirms rapid degradation of mecillinam in MOPSgluRDM

The rapid degradation which we inferred from our delay time bioassay was confirmed by direct measurement of mecillinam degradation using Raman spectroscopy. Raman spectra were measured for samples of mecillinam (at 3 mg/ml) in MOPSgluRDM, and for MOPSgluRDM alone, over a period of 60 hours (see Methods). Fig 4C shows part of the resulting difference spectra (shifted vertically for clarity). A peak at 1278 cm^−1^, which is believed to arise from the amidine link in the mecillinam molecule (see Fig 1 and Methods), clearly disappears over time. This is consistent with evidence that the primary degradation pathway for mecillinam is via hydrolysis of the amidine bond [16]. Fitting an exponential decay function to the peak height (relative to the background spectrum) at 1278 cm^−1^ as a function of time gives a half-life of 1.3 ± 0.4 hours (Fig 4D), consistent with our result from the delay time bioassay.

### Adjusting pH and temperature can increase mecillinam stability

To perform growth assays in the presence of mecillinam, one would like to find conditions under which mecillinam is more stable, while the bacterial growth rate is perturbed as little as possible. While low pH is known to increase the stability of mecillinam in aqueous solution [16], *E. coli* grows well only in the range pH 6.0 to 7.5 [41], and MOPS acts as a pH buffer only in the range pH 6.2 to 8.0 [22]. Therefore we repeated our delay-time bioassay in MOPSgluRDM medium at the lower pH of 6.5. This led to an increase in the half-life of mecillinam, to approximately 4 hours as measured with the delay-time bioassay (S4 Fig A), which was confirmed by Raman spectroscopy (S4 Fig D,E). Since this is still rather short on the timescale of a typical bacterial growth experiment, we sought to further optimise the conditions by decreasing the temperature. Therefore we repeated the delay time assay at 34°C, for pH 7.4 and pH 6.5. While a decrease in temperature to 34°C on its own did not significantly increase mecillinam stability (S4 Fig B), combining the temperature decrease with a pH of 6.5 led to a mecillinam half-life of more than six hours (S4 Fig C), while still maintaining an acceptable bacterial growth rate (Table 1). Further decreasing the temperature to 30°C resulted in a significantly lower bacterial growth rate (Table 1).

**Table 1.** Bacterial growth rates and antibiotic half-lives in MOPSgluRDM at varying temperature and pH. Growth rates were measured by applying a linear fit to the log of the optical density (the number of replicates used is indicated in the brackets). The antibiotic half-lives measured with the delay-time bioassay are also listed, for mecillinam (Mec.), cefotaxime (Ctx.) and aztreonam (Azt.).

| T (°C) | pH | Growth rate (/h) | MEC half-life (h) | CTX half-life (h) | AZT half-life (h) |
| --- | --- | --- | --- | --- | --- |
| 37 | 7.4 | $1.6 \pm 0.4$ (3) | $\sim 2$ | $> 6$ | $> 6$ |
| 37 | 6.5 | $1.2 \pm 0.1$ (10) | $\sim 4$ | | |
| 30 | 7.4 | $0.97 \pm 0.05$ (4) | | | |
| 34 | 7.4 | $1.3 \pm 0.1$ (3) | $\geq 2$ | | |
| 34 | 6.5 | $1.4 \pm 0.2$ (7) | $> 6$ | | |
| 37 | 7.0 (LB) | $1.5 \pm 0.2$ (4) | 4 – 6 | | |

### Mecillinam is somewhat more stable in LB medium than in MOPS

Luria-Bertani (LB) medium is an undefined growth medium that is widely used for bacterial growth experiments. We used the delay-time bioassay to determine the stability of mecillinam in LB medium at 37°C. The pH of the medium was measured to be 7.0 at the start of the experiment (although since LB does not contain a pH buffer we expect the pH to decrease during the experiment due to bacterial growth). Our results show that mecillinam has a half-life of 4-6 hours in LB at 37°C (S5 Fig). The fact that mecillinam appears to be more stable in LB than in our MOPSgluRDM experiments at pH 7.4 may arise from the somewhat lower pH of the LB medium [16]; another significant factor might be a lower concentration of transition metal ions, which have been found to destabilise *β*-lactams [32, 33].

### Aztreonam and cefotaxime have longer half-lives than mecillinam in MOPSgluRDM

To demonstrate the generality of the delay time assay, we used it to estimate degradation rates of two other *β*-lactam antibiotics in MOPSgluRDM at pH 7.4 and 37°C. Aztreonam is a monobactam that selectively targets the division transpeptidase PBP3 [42], while cefotaxime is a third-generation cephalosporin that targets multiple components of the cell division machinery [38]. Both these antibiotics have qualitatively similar effects on the growth dynamics of *E. coli*, with a concentration-dependent period of growth inhibition, followed by regrowth of the bacterial population (Fig 5A).

**Fig 5.**
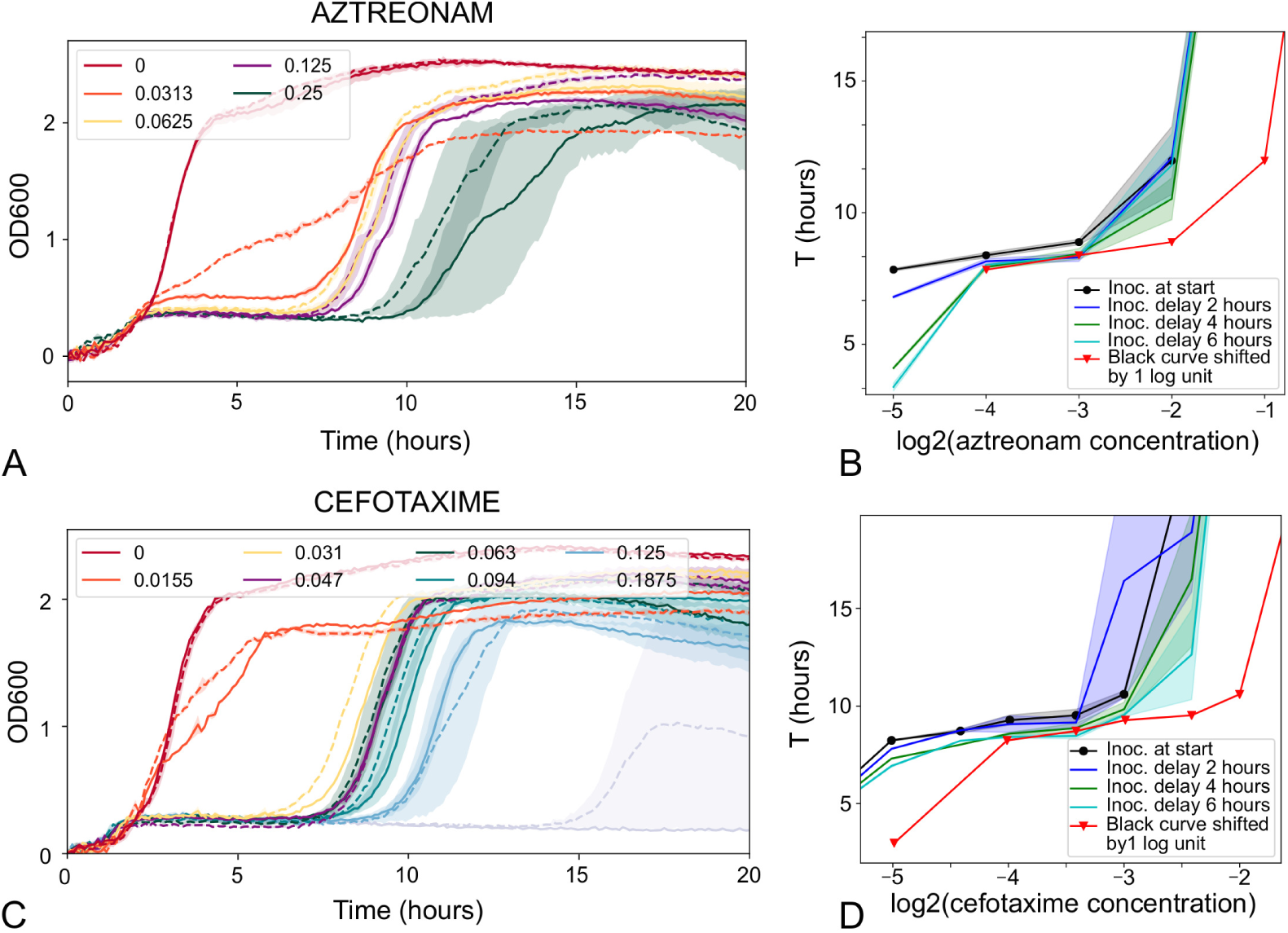
Aztreonam and cefotaxime have half-lives longer than 6 hours in MOPSgluRDM at pH 7.4 and 37°C. A,B: Delay-time bioassay for aztreonam. A: Time-shifted growth curves. Solid lines show results for immediate inoculation; dashed lines are for inoculation with a 4 hour delay. The aztreonam concentrations are listed in the legend with units of *µ*g/ml. B: Analysis (plots of time to reach OD threshold of 0.75). The red curve shows the black curve (immediate inoculation) shifted by 1 log_2_ unit. The blue curve corresponds to inoculation after a 2 hour delay, the green curve a 4 hour delay and the teal curve a 6 hour delay. Since the curves for delayed inoculation of 4h and 6h (green and teal curves) do not reach the red curve, the half-life is longer than 6 hours. C,D: Delay-time bioassay for cefotaxime. C: Time-shifted growth curves. Solid lines show results for immediate inoculation; dashed lines are for inoculation with a 2 hour delay. The cefotaxime concentrations are listed in the legend with units of *µ*g/ml. D: Analysis (plots of time to reach OD threshold of 0.75). The red curve shows the black curve (immediate inoculation) shifted by 1 log_2_ unit. The blue curve corresponds to inoculation after a 2 hour delay, the green curve a 4 hour delay and the teal curve a 6 hour delay. Since the curves for delayed inoculation of 4h and 6h (green and teal curves) do not reach the red curve, the half-life is longer than 6 hours.

For aztreonam, the delay time assay showed degradation over the course of the experiment, but with a half-life greater than six hours (Fig 5B). This is consistent with previous suggestions that monobactams are more stable than other *β*-lactams (except under alkaline conditions), possibly because of reduced strain on the *β*-lactam ring due to the absence of a secondary ring (Fig 1) [35].

For cefotaxime, the characteristic shape of the antibiotic-inhibited growth curve (i.e. a period of inhibition followed by regrowth) was only observed within a rather narrow concentration range (0.016 to 0.125*µ*g/ml). For smaller concentrations of cefotaxime, growth was not inhibited, while for larger concentrations, growth was completely inhibited (no regrowth was observed) (S6 Fig). Because the delay time bioassay relies on matching the shapes of growth curves for different antibiotic concentrations, the analysis could only be performed within this rather narrow concentration range where the growth curves were qualitatively similar (Fig 5C). Nevertheless, our analysis showed clearly that, although cefotaxime does degrade during our experiment, its degradation half-life is longer than 6 hours (Fig 5D - the curve for the 6 hour inoculation delay lies to the left of the zero-delay curve shifted by one log_2_ unit).

## Discussion

Microbiological growth assays, such as the MIC assay, play a central role in the diagnosis and treatment of bacterial infections. These assays usually assume that the antibiotic in question is stable over the timescale of the assay, typically 24 hours. Our results call this assumption into question, since we find that several *β*-lactam antibiotics degrade on faster timescales than 24 hours in widely used bacterial growth media, with mecillinam being especially prone to rapid degradation. This suggests that care is needed in the interpretation of MIC test results for *β*-lactam antibiotics. While rapid degradation of *β*-lactams in aqueous solution has been widely reported in the chemical literature, we believe that our study is the first to report this in standard microbiological growth media.

The delay-time bioassay which we develop here is easy to implement in a microbiological lab, requiring only a 96-well plate reader capable of shaking incubation and optical density measurement. The underlying principle of the assay is very simple: replicate wells containing antibiotic plus media are incubated in the plate reader for different amounts of time prior to inoculation, and the shape of the resulting bacterial growth curve is used to infer the effective concentration of the antibiotic in a given well at the time of inoculation. To simplify our analysis, we defined a threshold optical density and plotted the time taken to reach this threshold versus the logarithm of the (initial) antibiotic concentration: this provides a simple way to extract the degradation half-life. This analysis works for the three *β*-lactam antibiotics which we studied here, because of the ‘regrowth’ phenomenon: bacterial populations typically eventually reach a high density in these antibiotics, after an initial period of growth suppression that depends on the antibiotic concentration. For cefotaxime, where this characteristic growth curve was only observed within a narrow concentration range, the analysis was somewhat more challenging but the method nevertheless allowed us to place a lower bound on the degradation time.

For other classes of antibiotic, we would expect different characteristic forms for the growth curves, potentially making the definition of a threshold optical density more difficult. However, the general principle of matching growth curves for later inoculation times with those of reduced initial antibiotic concentrations still holds. Our assay also relies on the principle that replicate growth curves are expected to be very similar. This is the case for the *β*-lactams that we study here (Figs 4, 5) but variability between replicate growth curves might be an issue for some other antibiotics, at concentrations close to the MIC. In this case one might need to perform averages over larger numbers of replicates, inoculated at the same time.

The three antibiotics studied here, mecillinam, aztreonam and cefotaxime, represent different *β*-lactam sub-classes and show differing stability, with mecillinam being particularly unstable. However, we find that all of them degrade over timescales that are significantly shorter than a typical bacterial growth assay, suggesting that MIC and other assays should be interpreted carefully.

While our study focused on standard microbiological growth media (MOPS-based media and LB), it would also be interesting to measure antibiotic stability in media that are more typical of infections, such as urine or tissue culture medium. Recent work suggests that antibiotic susceptibility can be significantly altered in tissue culture medium compared to standard bacterial growth media [43]; it would be interesting to know if the same is true for antibiotic degradation. If *β*-lactam degradation is indeed rapid under conditions that are realistic of clinical infections, this could be a significant factor in determining antibiotic dosage protocols.

## Materials and methods

### Antibiotics

Antibiotic stock solutions were as follows: mecillinam (Sigma-Aldrich), 3 mg/ml in sterile water; cefotaxime (Fisher-Scientific), 10 mg/ml in sterile water; aztreonam (Cambridge Bioscience), 25 mg/ml in DMSO. Antibiotic solutions prepared by dilution of the stock solutions in water were filter sterilised using 0.22 *µ*m membrane filters (MillexGP).

### Bacterial strains

MIC (minimum inhibitory concentration) assays to determine the antibiotic concentrations of interest were performed using *Escherichia coli* strain MG1655. Bioassay measurements were performed using *E. coli* strain RJA002 [44], which is a derivative of MG1655 carrying a chromosomal yellow fluorescent protein (YFP)/chloramphenicol resistance reporter. The reporter is under the control of the constitutive *λ*P_*R*_ promoter, and was made by P1 transduction from strain MRR of Elowitz et al. [45]. Growth curves for RJA002 in the presence of the relevant *β*-lactam antibiotics were found to be indistinguishable from those obtained for the wild-type MG1655 strain (see supplementary material S3 Fig A and B).

### Microbiological growth media

Luria-Bertani (LB) medium was made in-house using components purchased from Fisher-Scientific, and its pH was 7.0. MOPS rich defined medium with glucose (referred to here as MOPSgluRDM) is a version of Neidhardt’s MOPS defined medium supplemented with nucleotides, amino acids and vitamins [22]. It was prepared in-house following the protocol developed by the *E. coli* Genome Project (University of Wisconsin) [46] to make EZ Rich Defined Medium (RDM). In this protocol, EZ RDM is assembled from four components (10x MOPS base medium, 0.132M K_2_HPO_4_, 10x ACGU (containing nucleotides) and 5x EZ supplement (containing amino acids and vitamins) - for details see below). Each component was made, sterilised and stored as single use aliquots. The complete medium was assembled just prior to use. Carbon source was also added at this stage. The in-house MOPS media was compared with commercially available MOPS media (Teknova), and found to support the same growth dynamics and doubling times for *E. coli* MG1655 in MOPSgluRDM (doubling time 34 ± 1 minutes) as well as in MOPSgluMIN (MOPS media without added amino acids, nucleotides or vitamins; doubling time 103 ± 2 minutes) (S1 Fig).

The MOPS base medium consisted of: MOPS buffer (40mM, pH 7.4 with KOH), 4mM Tricine (source of iron), 10*µ*M iron sulphate, 9.5mM ammonium chloride, 0.276mM potassium sulphate, 0.5*µ*M calcium chloride, 0.525mM magnesium chloride, 50mM sodium chloride, 29nM ammonium molybdate, 4*µ*M boric acid, 0.3*µ*M cobalt chloride, 96nM copper sulphate, 0.812*µ*M manganese chloride, 98nM zinc sulphate, 1.32mM dipotassium phosphate, sterile H2O and K_2_HPO_4_. The resulting MOPS growth medium has a buffering capacity from pH 6.2 to 8.0 [22] and was supplemented with glucose to a final w/v of 2%. The EZ supplement consists of the following amino acids (concentrations listed are those in the complete MOPSgluRDM media): 0.8mM alanine, 5.2mM arginine, 0.4mM asparagine, 0.4mM aspartic acid, 0.1mM cysteine, 0.6mM glutamic acid, 0.6mM glutamine, 0.8mM glycine, 0.2mM histidine, 0.4mM isoleucine, 0.8mM leucine, 0.04mM lysine, 0.2mM methionine, 0.4mM phenylalanine, 0.4mM proline, 10mM serine, 0.4mM threonine, 0.1mM tryptophan, 2mM tyrosine, 0.6mM valine. Vitamins are added to the EZ supplement: namely thiamine HCL, calcium pantothenate, p-aminobenzoic acid, p-hydroxybenzoic acid, and 2,3 –dihydroxybenzoic acid each at a final concentration of 10*µ*M. The ACGU supplement consists of 0.2mM adenine, 0.2mM cytosine, 0.2mM uracil, 0.2mM guanine.

In this work, where needed, MOPSgluRDM was adjusted to pH 6.5 by gradually adding 1M HCl. The final volume of 1M HCl added was 2-3% of the total volume.

### Raman spectroscopy

Raman spectra were measured for samples of mecillinam in MOPSgluRDM at pH 7.4 or 6.5, at a concentration of 3 mg/ml. Measurements were taken at various timepoints over several days. One to two independent spectra were obtained at each timepoint. Spectra were also measured for samples of MOPSgluRDM only; these were obtained as averages of three independent measurements. The Raman spectra were obtained using a high contrast Coderg triple spectrometer set at a resolution between five and six wavenumbers across the recorded spectral region. The excitation source was an argon ion laser operating at 488 nm with plasma lines removed by a prism spectrometer to produce approximately 1W pure laser light though the cell. The sample cuvette was fitted into a copper block with 4 holes at right angles to let the beam through and allow right-angle Raman collection (S2 Fig). The copper block was temperature controlled to better than 0.1°C with a Eurotherm controller in P.I.D. mode and maintained at 37°C. To reduce the time taken per scan, allowing better time resolution, spectra were recorded only in the range 0-1800cm^−1^. Following data acquisition, difference spectra were obtained by subtracting the (averaged) MOPSgluRDM spectra for a given timepoint from those of the mecillinam-MOPSgluRDM samples at the same timepoint, using Origin software. The resulting difference spectra were then cross-referenced with the standard mecillinam spectrum available in Biorad’s KnowItAll software. Using peak height relative to the background signal (10-20 wavennumbers away from the peak), peaks were identified which showed clear decay over time. These peaks were assigned using Biorad’s KnowItAll software, combined with a literature survey [47–55]. One peak of interest was determined (see Fig 4C): 1278 cm^−1^ (thought to be due to twisting of C-N-C [47], or twisting of the CH_2_ group [55])). The peak at 1278 cm^−1^ was identified as important as a Biorad’s KnowItAll database search of possible mecillinam degradation products (e.g. penicilloic acid [16]) demonstrated that these degradation products generally are not expected to have a peak at 1278 cm^−1^. In summary this peak is thought to be linked to the amidine bond found in mecillinam (Fig 1 A). The background signal used for the 1278 cm^−1^ peak was the average signal from 1261 - 1266 cm^−1^.

### Plate reader growth

Bacterial growth measurements were performed in a Fluostar Optima plate reader, with double orbital shaking at 600rpm, using Greiner flat-bottomed 96-well plates. Lids were used to reduce evaporation over the course of long experiments. The lids were taped onto the plate using small strips of autoclave tape on each side, allowing air to flow while preventing the production of plastic dust which occurs for shaken plates with unsecured lids. The optical density (OD) of each well in the plate was measured approximately every five minutes. The optical density data from the plate reader was analysed using custom Python code to determine growth rates and averages across replicates. A typical plate setup is shown in Fig 3. Growth rates were extracted from plots of the natural log of the optical density. A linear fit over a window of 10 datapoints (or 20 for very slow growth) was used. This window was moved along the log(OD) curve one datapoint at a time, with a linear fit determined at each step i.e. a gradient and residual was found for each window. The best-fit maximal growth rate is taken as the highest gradient with the lowest residual. This is necessary as a low gradient with a good linear fit (low residual) is likely to be from the stationary phase portion of the growth curve

### MIC assays

Minimum inhibitory concentrations (MIC) were measured via a micro-dilution protocol [56] using a Fluostar Optima plate reader in LB medium. For each antibiotic, the concentration range used in the MIC assay was determined from the MIC database on the EUCAST (European Committee on Antimicrobial Susceptibility Testing) website (Table 2). Each MIC assay was performed in duplicate columns of a 96-well plate, with each well containing 190 *µ*l of antibiotic-supplemented growth media. Within each column, the top row contained the highest concentration of antibiotic (generally four times the EUCAST MIC value), and the antibiotic concentration was diluted 2-fold down the column, with the bottom well containing no antibiotic. Each well was then inoculated with 10 *µ*l of *E. coli* MG1655 that were growing exponentially at OD∼0.2. After inoculation the plate was incubated for 22-24 hours at at 37°C with orbital shaking at 600 rpm (in the plate reader). The MIC was determined by visual inspection of the turbidity of the wells after incubation: the MIC was taken to be the lowest concentration that showed no turbidity in either of the duplicate wells [56]. If the antibiotic concentration range obtained using the EUCAST database did not result in an MIC value (i.e. if all wells showed growth), then the concentration range was shifted and the assay repeated until an MIC value was obtained. In each plate, at least one column contained only growth medium as a control for contamination. The MICs measured in this work are also listed in Table 2, and were generally slightly higher than those reported in the EUCAST database. This could be due to differences in measurement protocol e.g. inoculation size used (we note that EUCAST MIC data are aggregated from various sources which may use slightly different protocols).

**Table 2.** MIC values for the antibiotics used in this work.

| Antibiotic | EUCAST MIC range ($\mu\text{g/ml}$) | MIC as measured in this work ($\mu\text{g/ml}$) |
| --- | --- | --- |
| Cefotaxime | 0.016-0.25 | 0.25 |
| Mecillinam | 0.064-1 | 1.5 |
| Aztreonam | 0.03-0.25 | 0.5 |

### Delay-time bioassay

The delay-time bioassay measures antibiotic degradation rates by sequential bacterial inoculation of identically-prepared wells in a 96-well microplate, at regular time intervals. It is an adaptation of the MIC protocol that takes advantage of the fact that if an antibiotic is degrading over time we should observe a change in the growth dynamics between sequential inoculations. For example, if an antibiotic has a degradation half-life of two hours, and a bacterial inoculum is added two hours after the plate is filled with antibiotic, at what was initially a concentration of X *µ*g/ml, the culture should exhibit the growth dynamics normally associated with a concentration of X/2 *µ*g/ml.

In the delay-time bioassay, a 96-well plate is set up as in Fig 3: 6 columns of the plate are filled with 190 *µ*l of antibiotic-supplemented growth medium per well. Within each column, the antibiotic concentration is diluted 2-fold in successive rows, and the bottom row contains no antibiotic. The starting concentration is set at four times the measured MIC (Table 2), therefore the third row is at the MIC. At least one additional column contains growth medium only, as a control for contamination. At the start of the experiment, the first 2 columns are inoculated with 10*µ*l of a “starter” culture of *E. coli* (strain RJA002), in the exponential phase of growth at OD_600_ ∼0.2. Following inoculation, the OD_600_ of the inoculated wells is ∼0.01 and the final volume is 200 *µ*l. The plate is then incubated in the plate reader at 37° C, with 600rpm double orbital shaking, and OD measurements are performed at time intervals of ∼5 min. At the same time, the starter culture is diluted such that it is expected to again reach OD∼0.2 after a period of 2 hours. After the plate and starter culture have been incubated for 2 hours, the plate is removed from the plate reader, the next 2 columns are inoculated from the starter culture, the starter culture is again diluted, and incubation (with plate reader OD measurement) is continued. This process is repeated until all wells have been inoculated. The plate is then incubated in the plate reader, with OD measurement, for a further 16-18 hours.

Following data acquisition, a background subtraction is performed for each well, using the first OD reading (time zero) as a proxy for the background OD (note that the inoculum size is small enough to be below the plate reader’s detection threshold). Background-subtracted OD measurements for replicate wells are then averaged, and each set of data is time-shifted by its inoculation time, to produce a growth curve starting from the time of inoculation. This results in a set of 6 growth curves, corresponding to bacterial growth in medium that has been pre-incubated for 0, 2, 4, 6, 8 or 10 hours prior to inoculation (see Fig 4A). If the antibiotic does not degrade, these curves would be identical; antibiotic degradation will cause the curves to shift with the time of pre-incubation, since the antibiotic concentration experienced by the bacteria is lower than the initially added concentration.

To quantify the extent of any shift in the growth curves, we define a threshold OD value (OD_*t*_=0.75 in this work), and measure the time *T* after inoculation at which this threshold is reached. This time *T* increases with the antibiotic concentration, since the antibiotic inhibits bacterial growth. (Fig 4B). We plot a curve of *T* versus log_2_(antibiotic concentration), for each inoculation time. In this plot, degradation of the antibiotic results in a change in the effective antibiotic concentration experienced by the bacteria, and thus a shift to the right of the plot. The antibiotic half-life is the inoculation delay required to shift the curve by one unit on the logarithmic concentration axis (Fig 4B).

## Acknowledgments

RB was funded by an EPSRC DTA studentship. RJA and AD were funded by the European Research Council under ERC Consolidator grant 682237 EVOSTRUC; RJA was also funded by a Royal Society University Research Fellowship. We thank S. Lucas Black for making strain RJA002 and Meriem El Karoui and Bar lomiej Waclaw for helpful discussions.

